# Identifying a Distractor Produces Object-Based Inhibition in an Allocentric Reference Frame for Saccade Planning

**DOI:** 10.1101/2024.01.30.578028

**Authors:** Coleman E. Olenick, Heather Jordan, Mazyar Fallah

**Affiliations:** Department of Human Health and Nutritional Sciences, College of Biological Science, University of Guelph, Guelph, Ontario N1G 2W1, Canada; Canadian Action and Perception Network, Canada

## Abstract

We investigated whether distractor inhibition occurs relative to the target or fixation in a perceptual decision-making task using a purely saccadic response. Previous research has shown that during the process of discriminating a target from distractor, saccades made to a target deviate towards the distractor. Once discriminated, the distractor is inhibited, and trajectories deviate away from the distractor. Saccade deviation magnitudes provide a sensitive measure of target-distractor competition dependent on the distance between them. While saccades are planned in an egocentric reference frame (locations represented relative to fixation), object-based inhibition has been shown to occur in an allocentric reference frame (objects represented relative to each other independent of fixation). By varying the egocentric and allocentric distances of the target and distractor, we found that only egocentric distances contributed to saccade trajectories shifts towards the distractor during active decision-making. When the perceptual decision-making process was complete, and the distractor was inhibited, both ego- and allocentric distances independently contributed to saccade trajectory shifts away from the distractor. This is consistent with independent spatial and object-based inhibitory mechanisms. Therefore, we suggest that distractor inhibition is maintained in cortical visual areas with allocentric maps which then feeds into oculomotor areas for saccade planning.

## Introduction

When I view a tabletop full of coffee cups, I can locate my cup relative to myself in a viewpoint-dependent, egocentric reference frame. When I want to reach for my cup, there is considerable evidence that both my eye- and hand-movements are programmed using an ego-centric spatial map of the scene. However, if I want to avoid knocking over my colleagues’ cups, I need to direct my actions towards my cup while considering the locations of the non-target cups. This requires an allocentric (object-to-object) spatial map of the scene, where objects are represented in a viewpoint-independent reference frame. There is converging evidence from behavioral, electrophysiological, and imaging studies that we use both viewpoint-dependent egocentric, and viewpoint-independent allocentric reference frames in spatial cognitive tasks using reaching^[1][2][3][4]^.

When there are multiple objects in the visual scene, attending to and acting on, relevant stimuli may require the suppression of irrelevant stimuli competing for attentional selection. Yet there is no clear delineation about the reference frames supporting this suppression. In models of attention based on visual processing for example, the biased competition and selective tuning models, the suppression of irrelevant stimuli changes in strength based on the proximity to an attended stimulus^[5][6]^. However, oculomotor models of spatial attention invoke regions encoding relevant and irrelevant object locations in retinotopic coordinates, notably the superior colliculus (SC) and frontal eye fields (FEF)^[7][8][9]^.

The receptive fields of oculomotor regions also receive inputs in non-retinotopic reference frames, for example eye position-based gain modulations of the FEF^[10][11]^. There is also extensive work on the study of gain fields in parietal regions representing multiple reference frames^[12][13]^ that feed into the FEF and SC. While these inputs provide a source of non-retinotopic information, their effects are largely modelled as multiplicative gain factors on an underlying retinotopic map giving rise to the oculomotor plans.

Recently, saccadic eye movement trajectories have been shown to reflect the continuous competition resolution between objects^[14]^. Specifically, at short object processing times, before the target object has been differentiated from a distractor and decision making is still actively taking place, saccades deviate towards the distracting object. However, once the target and distractor are differentiated after greater processing time, and decision making has completed, the saccades deviate away from the distractor. The directional deviation of a saccade can be used to differentiate between a distractor or location that has been inhibited and one that has not, for a given single saccade^[15–26]^. Saccades are planned in an egocentric reference frame on a retinotopic map of space^[27]^ and activity on this map has been shown to cause saccade trajectory deviations^[28–30]^. The deviation towards a distractor is due to increased activity at the location of the distractor, whereas suppression at the distractor location leads to deviation away^[25][29–30]^. Because saccade planning is egocentric^[31–33]^, it is assumed that the suppression of distractors would also be egocentric.

However, behavioral studies in humans have shown that object-based suppression can also occur in allocentric reference frames. For example, object-based inhibition of return (IOR) has been shown to occur in an allocentric reference frame^[34–37]^. Both trajectory deviations and IOR can occur under the same conditions but do so on different timescales^[38]^. The complex objects used by Giuricich, and colleagues^[14]^ required late-stage visual processing to discriminate target from distractor^[39–40]^. These late-stage ventral stream areas (e.g. TE, TEO, IT cortex) support object representations in allocentric reference frames^[41]^, likely resulting from large, bilateral receptive fields with position and size invariance^[41–43]^. So, if saccade planning occurs in an egocentric reference frame, but the resolution of object competition is dependent on regions with allocentric representations and both involve inhibition of a distractor, what is the reference frame of the resulting distractor suppression?

In this study we independently varied the egocentric and allocentric distance of a target and distractor to investigate the reference frame of distractor suppression. The egocentric distance of the target and distractor is their eccentricity, while the allocentric distance describes the distance between the target and distractor or target-distractor distance (T-D distance), which is independent of the viewpoint or eye position. Varying the eccentricity of the target and distractor while keeping the angle between them constant, leads to a range of T-D distances. Using saccade trajectories to infer the state of decision-making, we identified when the distractor has been suppressed through the resulting deviation away from the distractor. The magnitude of the saccade deviation is associated with the strength of the distractor-related activity as has been demonstrated through both recording and stimulation studies of the SC^[28–29]^ and FEF^[30]^. The amount of deviation at different egocentric and allocentric distances will reflect the reference frame of distractor activation and suppression. As the oculomotor system encodes saccades to a target in an egocentric reference frame, we expect to find egocentric reference frame effects on saccade trajectories both during active competition before the decision is made and after, when the distractor is suppressed. However, if distractor suppression is dependent on object-based inhibition developing in cortical visual areas, once the competition is resolved, saccade trajectories should also be shifted based on T-D distance in an allocentric reference frame.

## Methods

### Participants

Forty naïve participants (8 males; age: 18-27, mean = 19.6 years old) were recruited for the current experiment. All participants had normal or corrected to normal vision, gave written informed consent, and were compensated with $15 CAD for their time. Ethics approval of the experiment was obtained from the Research Ethics Board at the University of Guelph.

### Stimuli and Apparatus

Participants were seated in a dimly lit room 57cm away from a 21” CRT monitor (60Hz, 1024×768), with movement reduced by use of a chin and head rest. Stimulus presentation and data collection was controlled by Presentation software (Neurobehavioral Systems, Inc., Berkeley, CA), and participant responses were made using a response box (RB-540, Cedrus Corp., San Pedro, CA) and eye movements. The eye movements of participants were recorded from their right eye using an Eyelink II infrared eye tracker (500Hz; SR Research, ON, Canada).

The target and distractor stimuli were white (CIExy = [.29, 0.30]; luminance = 126.02cd/m^2^) on a black (CIExy = [.27, 0.26]; luminance = 0.20cd/m^2^) background and were randomly selected from a set previously used^[14][25][39]^ (Figure 1a). Each stimulus consisted of 6- or 7-line segments (1° × 0.08°). The stimuli resemble novel alphanumeric characters, requiring similar processing in late-stage ventral stream areas (e.g. TE, TEO, IT cortex)^[40][44]^.

**Figure 1:**
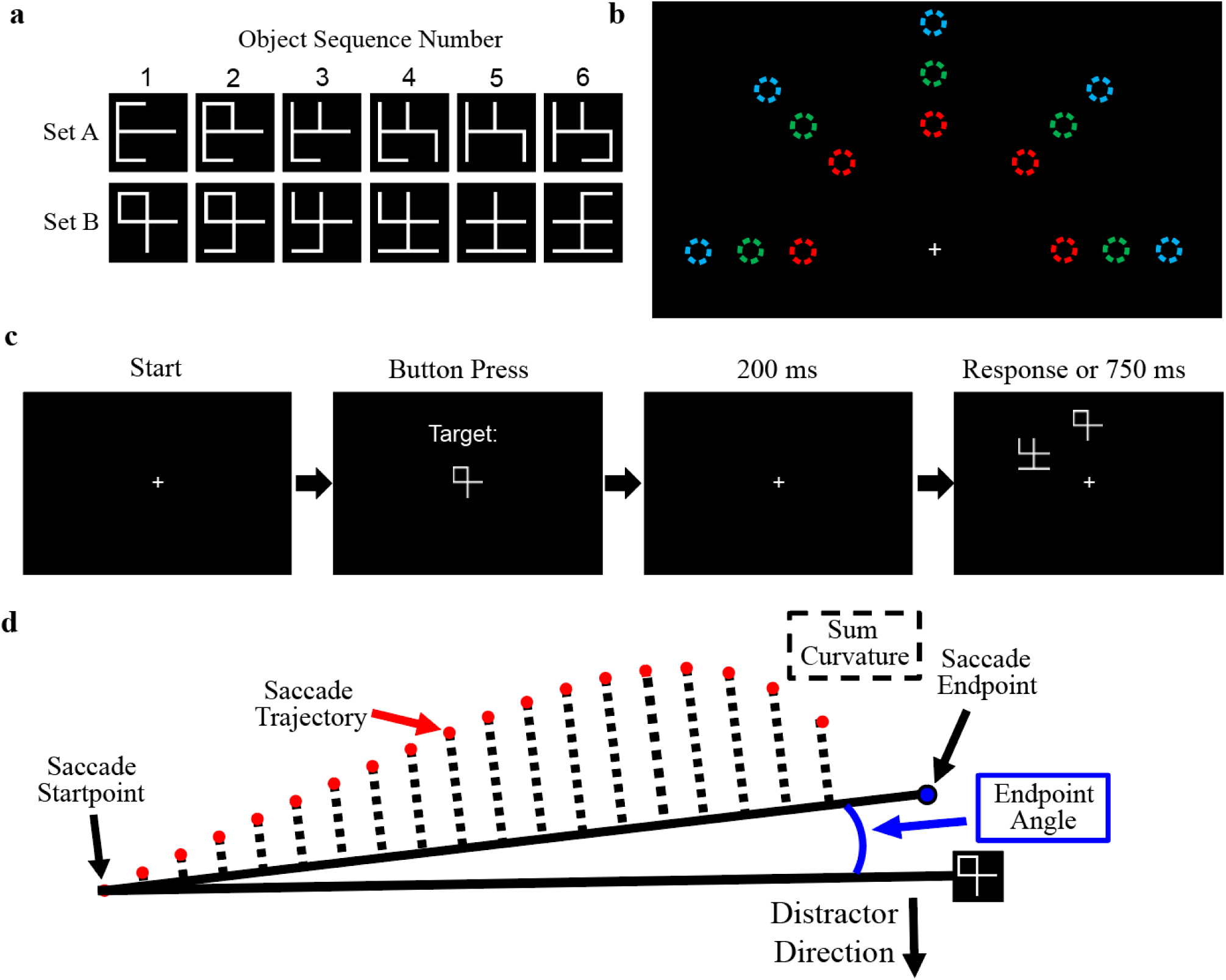
Experimental Overview. A) The stimuli were selected from 2 sets of 6 objects, each 2° X 2°. The target and distractor on a trial were drawn from the same set, with each object varying by one line addition or removal, relative to adjacently numbered objects in the set. The objects were designed to avoid resemblance to any English characters or numbers to ensure their novelty to participants. B) The objects were presented at 15 possible locations at 3 eccentricities (depicted in color for clarity in the figure only: 7 [red], 10 [green], and 13 [blue] dva from fixation), and 5 branches separated by 45° in the upper half of the display. The target and distractor were always objects from the same set and presented along adjacent branches. C) Trials began with participants fixating a central cross, which was replaced with a preview of the target for that trial. After participants pressed a button, they were required to again fixate a cross for 200ms before the target and distractor appeared simultaneously. The objects remained on screen until either a saccade was made to the target or distractor or 750ms had elapsed. Trials without a response in 750ms were presented again at a later point in the block, and there were some trials where only the target was presented to establish a saccade baseline. D) The sum curvature of a saccade was calculated as the area under the curve between the actual saccade trajectory and a path directly between its start and end points. The endpoint angle of a saccade was calculated as the polar angle between the saccade endpoint and the center of the target object. This example saccade is showing deviation away from the distractor direction and would be negative. See methods for more detail.

### Procedure

Participants completed a delayed match to sample task, making saccades to a previously shown target presented with a distractor (Figure 1c). Eye position was calibrated at the beginning of the experiment, at the start of each block, and additionally if necessary. Each trial began with a fixation cross (0.4° X 0.4°; CIExy = [.x=29, y=0.30]; 126.02cd/m^2^) at the center of the screen. Following 200 ms fixation, the target object was presented at the centre of the display and remained until the participant pressed a designated button on a response box. The target object was replaced by a fixation cross and the participant had to maintain fixation within a 2X2 dva square window centered on the cross for 200 ms or the trial was terminated. The fixation marker remained on the display while the target and a distractor object appeared simultaneously for 750 ms or until the participant made a saccade to one of the objects. Participants received a written error message and audible tone if they failed to make a saccade or made a saccade to the distractor object; error trials were presented later in within the same block.

There were 15 potential positions for the objects, organized into 5 different branches separated by 45°, and placed at egocentric eccentricities of 7, 10, or 13 dva (Figure 1b). The various eccentricities for the target and distractor yield a range of allocentric distances between them (T-D distance), ranging from 5.36 dva to 9.95 dva (5.36, 7.07, 7.65, 9.22, 9.45, and 9.95 dva). The target and distractor were always 45° apart, an angular distance previously investigated that has been shown to provide reliable saccade trajectory deviations^[14]^. Within each block, each combination of target and distractor eccentricities appeared with equal probability (3x3 combinations) and were evenly distributed across the array of possible positions (Figure 1b), resulting in 72 distinct target-distractor pairings (trials), pseudorandomly interleaved across the block. Note that all objects were presented on or above the horizontal meridian and thus when the target or distractor appeared on a horizontal branch, the other object necessarily appeared 45° above the meridian. However, this was not predictive as an object appearing on the horizontal meridian had equal probability of being the target or the distractor across the block of trials.

Additionally, each block contained target-only trials at each of the 15 object positions, acting as a baseline, producing a total block size of 87 trials. The objects used as target and distractor were always from the same set of the two sets used in the experiment (Figure 1a) but varied in their similarity to one another. This range of stimuli were used to increase the complexity of the task minimizing perceptual learning. Each participant completed 5 experimental blocks (435 total trials) and a 10-trial practice block presented at the beginning of the experiment that was not included in analysis.

### Data Analysis

The investigation of the effects of eccentricity made use of multiple regression across target and distractor eccentricities. To differentiate an allocentric reference frame from an egocentric reference frame, we a priori determined to investigate both trials with equal target and distractor eccentricity, and trials with equal T-D distance but flipped target and distractor eccentricities (e.g. target at 7 dva/distractor at 13 dva vs target at 13 dva/distractor at 7dva). All multiple comparisons tests and regression analyses have been subject to Benjamini-Hochberg corrections for multiple comparisons

#### Saccade Detection

Saccades were detected using in-house MATLAB (Mathworks, Natick, MA) scripts following pre-determined criteria; an eye movement with a velocity greater than 20°/s for at least 8 ms with a peak velocity greater than 50°/s. Saccades were evaluated manually to remove trials where the first saccade was not made to the target or distractor (9.59%). The saccades not made to the target or distractor were a mix of different outcomes: short saccades where the intended target of a saccade could not be determined, saccades that were initially made in the direction of one of the two objects but not reaching it before making a second saccade (a double-step saccade), and saccades that were partially occluded by an eye blink. Trials with short (amplitude < 1° [0.65%]) or express saccades (Saccade Reaction Time (SRT) < 100 ms [1.97%]) or where no eye position data was recorded (3.64%) were excluded from further analysis. Overall, 84.15% of recorded trials were included for analysis.

#### Accuracy

Accuracy was calculated for each participant as the proportion of trials that a saccade was made to the target rather than the distractor. A 4X4 dva box around the target or distractor was used to determine which object was selected for a given saccade. All participants’ performance exceeded chance (binomial distribution with p = 0.5 [50%]) with 65.2 ± 8.9% correct. *Saccade Trajectory Metrics*: To quantify the deviation of a saccade from a straight path, the sum curvature and endpoint angle for each saccade was calculated. The sum curvature provides a measure of the strength of target and distractor representations in the cortical saccade planning process^[17–18][30][45]^. Sum curvature was calculated as the total deviation of a saccade from a straight line between its start point and endpoint (reference line). That is, the magnitude of deviation from the reference line and direction (sign) of each eye position sample for a saccade were summed to generate a single metric for each saccade. The endpoint angle of a saccade is suggested to result from the outcome of a subcortical vector average of saccade plans to both the target and distractor^[44–45]^. The endpoint angle was calculated as the polar angle (in degrees) between the reference line and a straight path from the saccade start point to the center of the target. The reference line was translated prior to computation such that the start of each saccade was on the cartesian origin to account for small variations in initial fixation location. Both metrics were normalized by subtracting the mean values from target only trials and corrected such that positive values reflect deviations towards the distractor and negative values reflect deviations away from the distractor.

#### Processing Time Division

Following Giuricich and colleagues^[14]^, trajectory metrics were used to separate saccadic responses made between actively competing and inhibited distractor stages as these different processes may not rely on the same reference frame(s). When two items are potential saccade targets, a saccade made to one has its trajectory shifted towards the other. As an actively competing distractor in a perceptual discrimination is a potential saccade target, this results in saccade trajectories made to the target shifting towards the distractor^[15–18][21–24][29–30]^. In contrast, with enough time the discrimination decision-making is completed, wherein the target is identified and the distractor is inhibited. An inhibited distractor results in saccade trajectories shifting away from it^[19–20][25–26][29–30]^. The change in saccade deviation direction over time has been used to separate saccades executed at different stages of visual processing ^[14–26][29–30][45]^. To determine the range of saccade latencies falling within each stage (actively competing vs inhibited distractor), sum curvature was organized by saccade latency and a 10 ms width sliding window was used to create a series of binned saccades. Within each of these bins, a two-tailed t-test against 0 was performed to determine whether there was significant overall curvature towards or away from the distractor at each saccade latency (Figure 2b).

**Figure 2.**
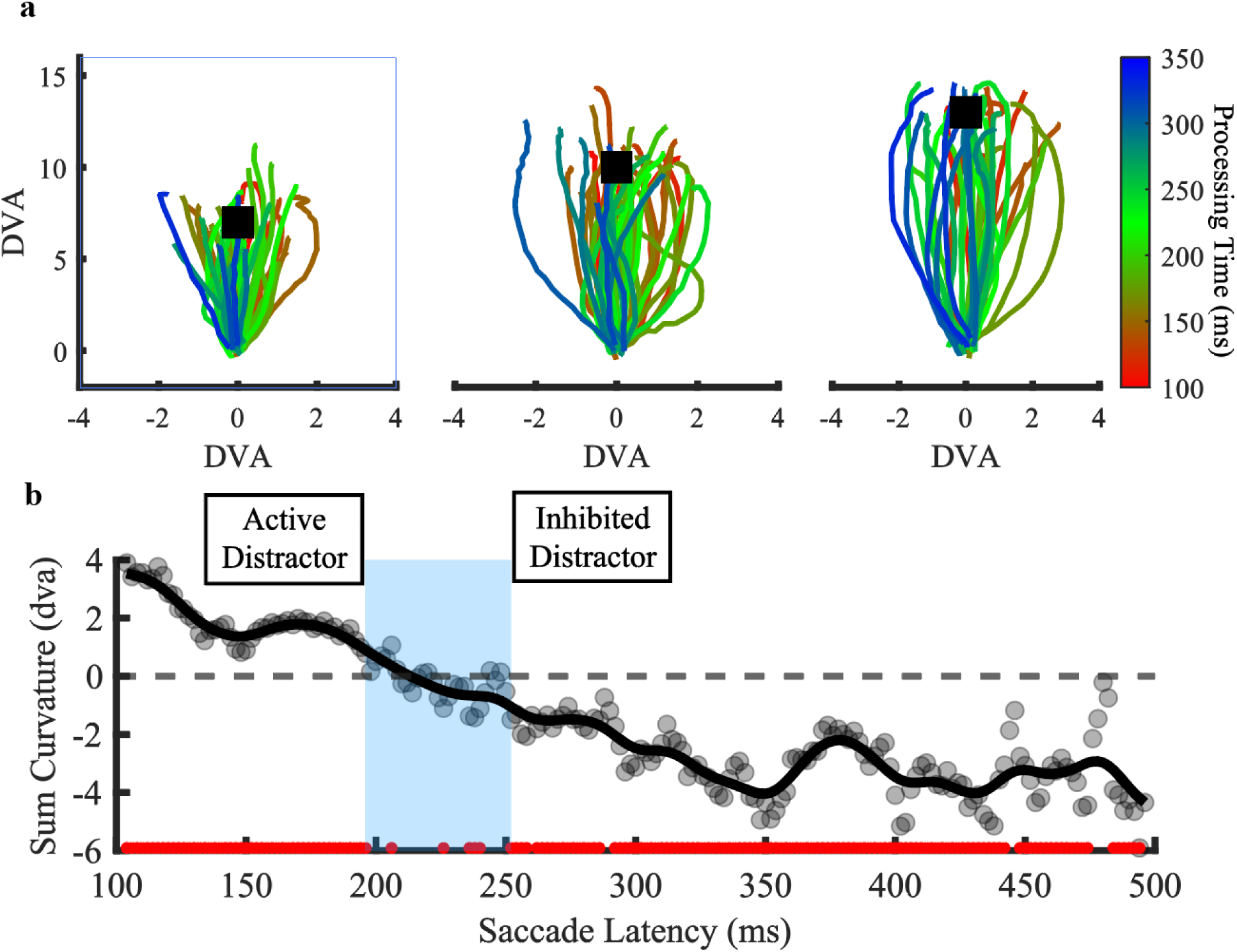
Raw Saccade Trajectories and Deviation Direction Over Processing Time. A) 80 random saccade traces to targets at 7, 10, and 13 dva (from left to right, respectively), rectified to a distractor appearing on the right, but uncorrected for saccade start position, are plotted. Black squares represent the center of the intended targets (note that the actual targets had a width of 2 dva), and the traces are color-coded by object processing time; red and blue reflecting near 100ms and 350ms respectively. B) The average sum curvature of saccades across all participants has been plotted over saccade latency. A 10 ms sliding window over saccade latency was used to group saccades and the average sum curvature within each bin is represented by the shaded grey points, with the solid line reflecting a 10 ms moving average over the shaded points. Positive values indicate deviation towards the distractor, where the distractor is still actively competing with the target, while negative values indicate deviations away after the distractor has been inhibited. The red points along the bottom represent the center of saccade latency bins where saccades showed significant curvature towards or away from the distractor (different from 0). The blue shaded region represents a transition between an active and inhibited distractor that has been omitted from further analysis.

## Results

Prior to analysis of the task performance and saccade trajectory metrics, the saccades were divided into two categories: those in which the distractor was actively competing with the target and those after it had been inhibited. This was accomplished by examining the saccadic sum curvature over object processing time (Figure 2b). By using a sliding 10 ms window to group saccades by object processing time and performing t-tests against 0 on sum curvature within each window, we identified time ranges with consistent significant deviation towards the actively competing distractor (100-196 ms) or away from the inhibited distractor (252-750 ms). In total, there were 10407 saccades outside the transition zone; 62.5% at short processing time and 37.5% at long processing times. These ranges are consistent with prior studies^[14][25][39][46]^. Saccades whose latencies fell in the transition zone between these 2 ranges were excluded from further analysis. It is important to note that not all participants had data in all combinations of experimental conditions and saccadic response times, making the use of repeated measures ANOVA impractical. Our analysis instead uses mixed-effects regression models as a well-established alternative^[47–49]^ for the purposes of the current study.

### Accuracy

In an egocentric reference frame, increasing the distance between fixation and the target would be expected to have the opposite effect as increasing the distance between fixation and the distractor, due to the central bias in attention (where attention and visual processing is stronger closer to fixation and drops off with distance) as shown by other visual search paradigms^[50–52]^. The central bias predicts that as the distance between the target and fixation increases, accuracy decreases, but as the distance between distractor and fixation increases, accuracy improves. To investigate the relationship between an egocentric reference frame and accuracy over processing times, a multiple linear regression with target and distractor eccentricity, and processing time (short and long) as factors was performed. The overall regression was found to be highly significant (R^2^ = 0.641, F(7,680) = 173.2, p < 6.4X10^-15^). Increasing target eccentricity was found to decrease accuracy, while increasing distractor eccentricity increased accuracy (Figure 3). The effect of both target and distractor eccentricity significantly decreased from short to long processing time. There was a significant interaction between target and distractor eccentricity that did not change with processing time, suggesting a relationship between object positions beyond their individual eccentricities, that will be explored in the next section. Finally, there was no change in accuracy across processing times independent from the other relationships mentioned above. However, a Wilcoxon signed rank test across each participant revealed a significant increase in accuracy at longer processing times (p = 3.19X10^-15^).

**Figure 3:**
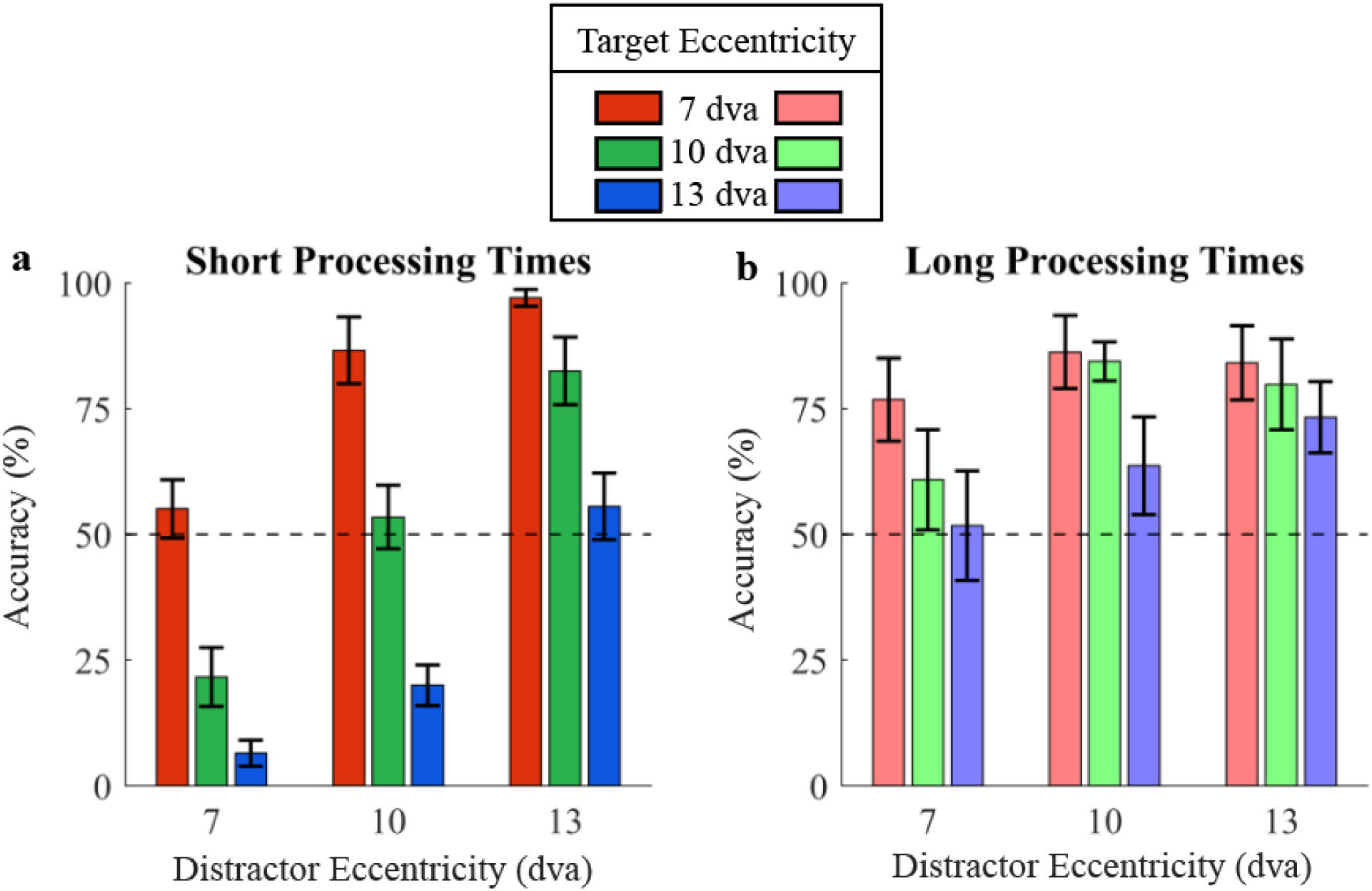
Accuracy by Target and Distractor Eccentricity. The average accuracy plotted over target and distractor eccentricity. A) Accuracy at short processing times while competition between target and distractor is still active. B) Accuracy at long processing times when competition has been resolved and the distractor had been inhibited. Error bars represent mean ± 95% confidence intervals. The bars and lines are colored according to the target eccentricity of 7 dva (red), 10 dva (green), and 13 dva (blue).

To identify any effects of an allocentric reference frame on accuracy, a multiple linear regression with processing time and target-distractor distance (T-D distance) was performed on the subset of trials in which the target eccentricity and distractor eccentricity were equal, i.e. equal egocentric distances (Table 1; Figure 4 B, C). When the target and distractor were both presented at an eccentricity of 7, 10, or 13 dva, the T-D distances were 5.36, 7.65, and 9.95 dva, respectively. The regression across T-D distance was significant overall, and robust across participants (R^2^: 0.332, F(3,224) = 37.14, p < 2.13X10^-15^). While overall accuracy increased significantly from short to long processing time (from regression: p = 0.0156), T-D distance had no influence on accuracy at short processing times. Critically, accuracy varied by T-D distance at long processing times, when the distractor is inhibited, decreasing significantly as the T-D distance increased.

**Figure 4:**
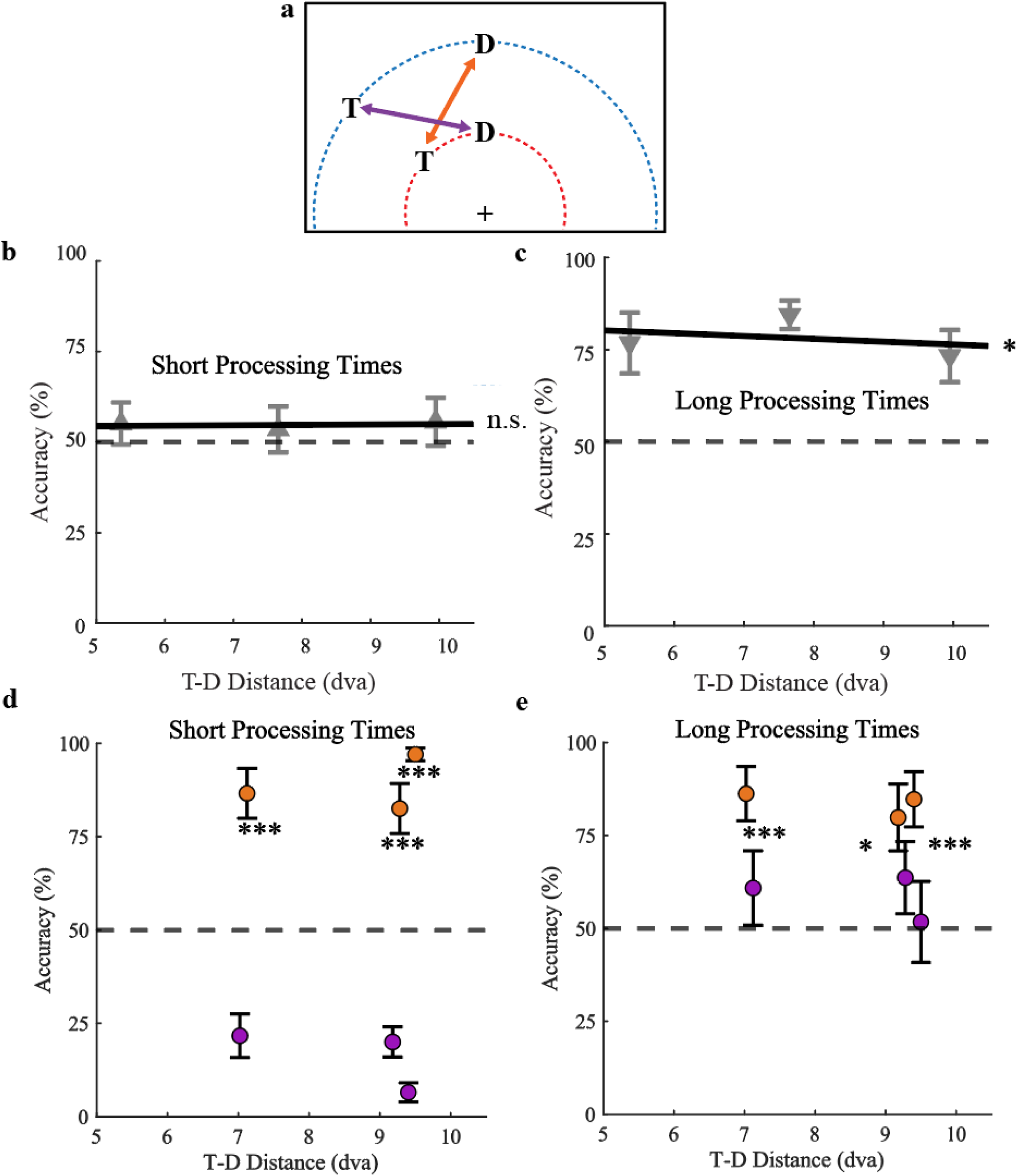
Accuracy by Target-Distractor (T-D) Distance Pairs. A) Example of target and distractor positions that have an equal distance between them (T-D distance) but either the target is closer to fixation (orange), or the distractor is closer to fixation (purple). Accuracy for targets and distractors presented at the same eccentricity but varying T-D distances plotted for B) short processing times and C) long processing times. The average accuracy plotted over T-D distance pairs for D) short processing times, and E) long processing times. Error bars represent mean ± 95% confidence intervals. Note that the pairs of orange and purple dots in panels B & C are offset slightly for visibility and that the actual T-D distances for each target closer and distractor closer pair are: 7.07, 9.22, and 9.45 dva from left to right. Pairwise comparisons between the allocentric distance pairs are indicated by n.s: not significant, *: p < 0.05 ***: p < 0.001.

**Table 1:** Regression Coefficients, and p-values for Accuracy and Trajectory Metrics Over Processing Time. Each cell consists of the regression coefficient and its p-value: *: p < 0.05, **: p < 0.01, ***: p < 0.001, n.s.: not significant. Note: the values reported for accuracy regressions were determined through separate linear regressions for short and long processing time after differences had been established. t is used to indicate coefficients that were not significantly different between processing times. The T-D distance regression was performed only on the points where target and distractor eccentricities were equal. The eccentricity interaction reflects a changing effect for the increase of target or distractor eccentricity as the eccentricity of the other object increases.

|  | Regression Predictor | Accuracy | Sum Curvature | Endpoint Angle |
| --- | --- | --- | --- | --- |
| Short Processing Time | Target Eccentricity | -0.139<br>$< 2 \times 10^{-16}$ *** | +1.040<br>$1.1 \times 10^{-4}$ *** | -1.392<br>$7.2 \times 10^{-5}$ *** |
| | Distractor Eccentricity | +0.035<br>$5.5 \times 10^{-3}$ ** | +0.039<br>0.8494 n.s. | -2.234<br>$< 2 \times 10^{-16}$ *** |
| | Eccentricity Interaction | +0.005<br>$3.11 \times 10^{-5}$ *** | -0.036<br>0.1282 n.s. | 0.153<br>$5.5 \times 10^{-7}$ *** |
| | T-D Distance | +0.004<br>0.521 n.s. | +0.446<br>$1.6 \times 10^{-4}$ *** | -0.915<br>$5.3 \times 10^{-12}$ *** |
| Long Processing Time | Target Eccentricity | -0.076<br>4.2x10<sup>-4</sup> *** | +0.168<br>0.4668 n.s. | -0.316<br>0.1061 n.s. |
|  | Distractor Eccentricity | -0.015<br>0.4843 n.s. | +0.261<sup>†</sup><br>0.2407 n.s. | -0.442<br>0.0206 * |
|  | Eccentricity Interaction | 0.004<sup>†</sup><br>0.0671 n.s. | -0.047<sup>†</sup><br>0.0313 n.s. | +0.040<br>0.0338 * |
|  | T-D Distance | -0.023<br>0.0104 * | -0.669<br>2.3x10<sup>-8</sup> *** | +0.074<br>0.511 n.s. |

The other method we employed to differentiate egocentric and allocentric coding was comparing conditions where the T-D distance between the target and distractor was identical, but their eccentricities varied Using paired t-tests (Figure 4 D, E), we see significant differences between eccentricities with the target or distractor closer to fixation at both processing times (all p < 1.933X10^-3^). However, the differences between the target closer and distractor closer conditions decreased substantially, indicating a reduction in the influence of an egocentric reference frame. Taken together, the reduction in center bias and variation in accuracy across T-D distance shown in the significant regression in the previous section at long processing times indicate that accuracy is driven by object distance in both egocentric and allocentric reference frames.

### Saccade Trajectory Deviations

To better understand the reference frames of target and distractor representations, we next investigated their effects on saccade trajectory metrics, which provide a continuous and separable measure of distractor activity and suppression. Separate multiple linear regressions were performed on the saccade curvature and endpoint angle for short and long processing times with target and distractor eccentricity as predictors (see Table 1; Figure 5).

**Figure 5:**
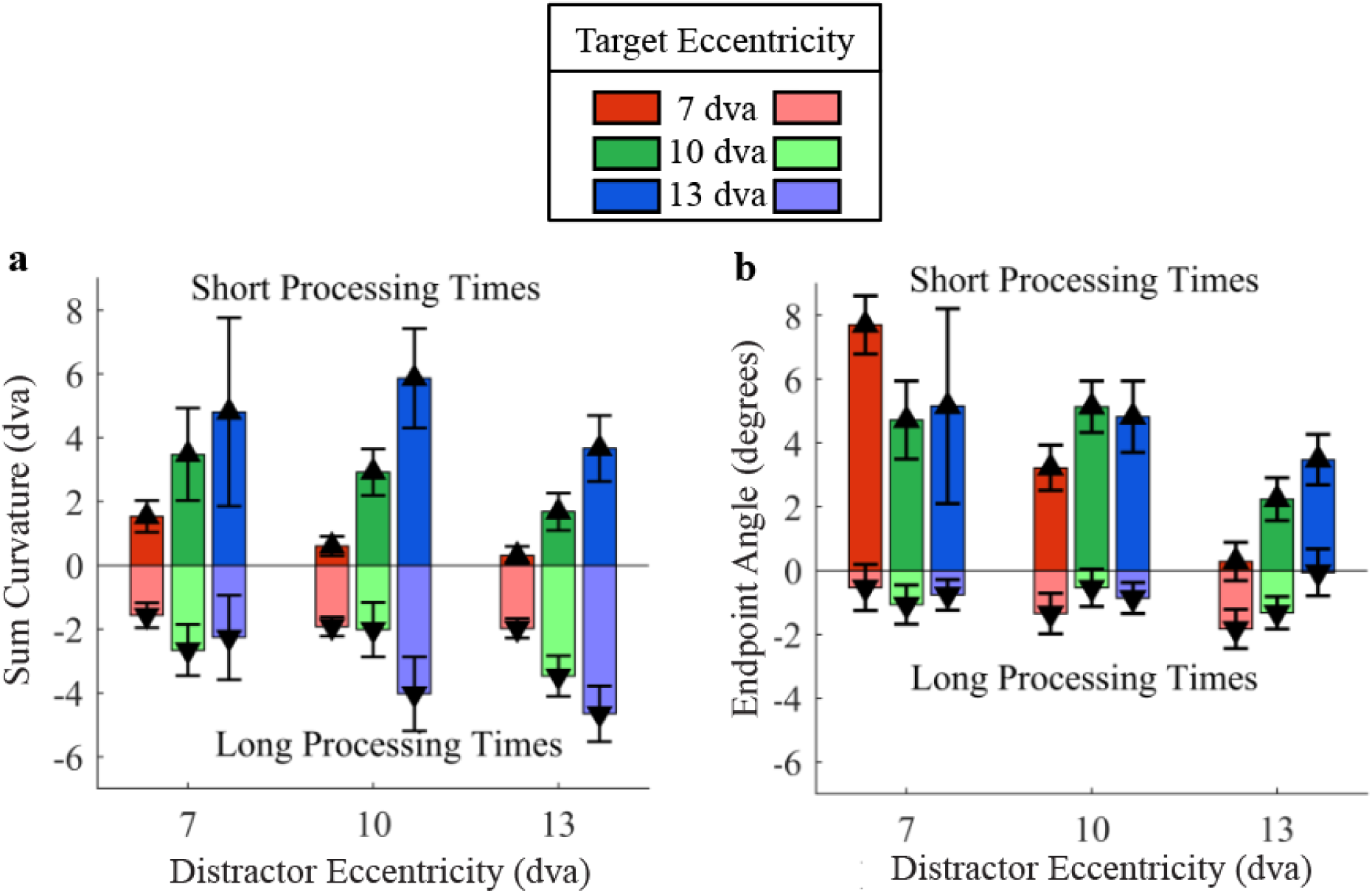
Saccade Trajectory Metrics by Target and Distractor Eccentricity. The average A) sum curvature and B) endpoint angle plotted over target and distractor eccentricity. Error bars represent mean ± 95% confidence intervals. Values above 0 reflect saccade deviations towards a distractor while the decision-making process is active. Values below 0 represent deviations away from the distractor, after object competition has been resolved and the distractor has been suppressed. The bars and lines are colored according to the target eccentricity of 7 dva (red), 10 dva (green), and 13 dva (blue).

At *short processing* time, both regressions were significant overall (Sum Curvature: R^2^ = 0.0449, F(3,3658) = 57.39, p < 9.11X10^-16^ | Endpoint Angle: R^2^ = 0.0598, F(3,3658) = 77.53, p < 7.98X10^-16^). At short processing time, where saccades deviate towards the distractor, only increasing target eccentricity led to greater saccade curvatures, consistent with an egocentric target representation whose strength diminishes with eccentricity. In contrast, the endpoint angle decreased as the eccentricity of the target and the eccentricity of the distractor increased, again consistent with an egocentric representation. A significant interaction effect between target and distractor eccentricity on endpoint angle indicates that the relative effect of the eccentricity of an object changes with the eccentricity of the other object, and may be suggestive of an allocentric effect and will be followed-up on in later analyses.

At *long processing* times, regressions for sum curvature and endpoint angle were both small but significant (Sum Curvature: R^2^ = 0.018, F(3,3176) = 19.43, p = 5.8X10^-12^ | Endpoint Angle: R^2^ = 0.0025, F(3,2492) = 3.045, p = 0.0051). Changes to the target eccentricity had no effect on the sum curvature or endpoint angle. The distractor eccentricity was correlated with the endpoint angle, but not the sum curvature. Interestingly, the interaction between target and distractor eccentricity was significantly correlated with both trajectory metrics, again suggesting target-distractor distance effects in an allocentric reference frame. Together, these results show a decreased strength of egocentric object representations feeding target-distractor competition in saccade planning after the decision-making process has been resolved.

To investigate the influence of an allocentric reference frame, linear regressions were performed on T-D distance for both saccade metrics when the target and distractor eccentricity were equal (Figure 6 A, B). In addition, pairwise t-tests were used to compare conditions where the T-D distance of the objects was equal, but their eccentricities varied (Figure 6 C-F). At *short processing* times, T-D distance had a significant effect on both the sum curvature (R^2^ = 0.0115, F(1,1231) = 14.37, p = 2.9X10^-4^), and endpoint angle (R^2^ = 0.0379, F(1,1231) = 48.53, p = 1.54X10^-11^). However, pairwise comparisons across equal T-D distance pairs also found significant differences for both metrics (all p < 0.0432). Together these findings suggest that at short processing times saccade trajectories are driven by egocentric rather than allocentric representations.

**Figure 6:**
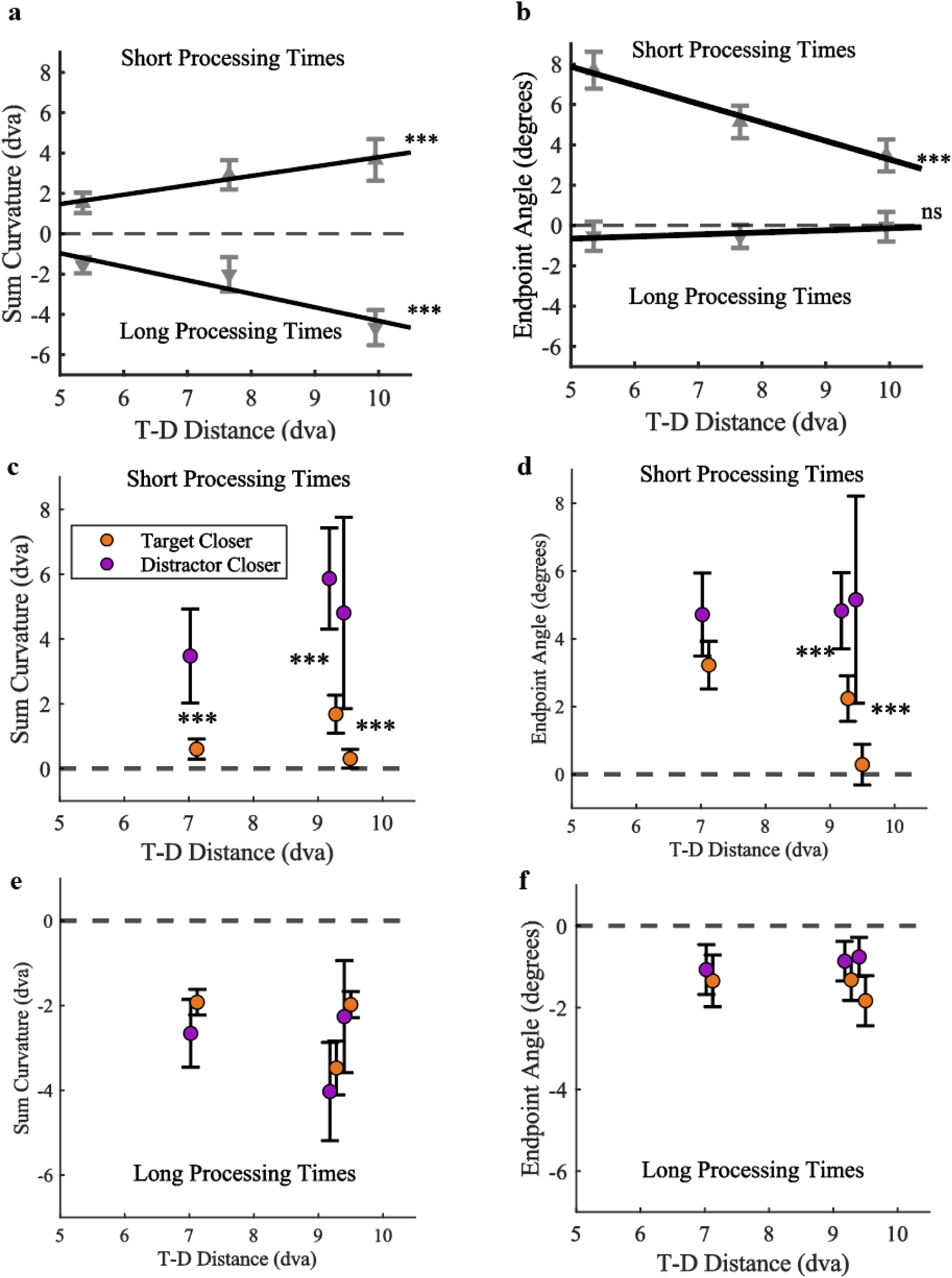
Target-Distractor (T-D) Distance. A & B: The average sum curvature and endpoint angle plotted over T-D distance when the target and distractor have equal eccentricity. Values above 0 reflect saccade deviations towards a distractor while the decision-making process is active. Values below 0 represent deviations away from the distractor, after object competition has been resolved and the distractor has been suppressed. C-F: The saccade metrics plotted over T-D distance pairs with the same conventions as in Figure 3, where orange reflects the conditions with the target presented closer to fixation and purple reflects trials with the distractor closer to fixation. Error bars represent mean ± 95% confidence intervals. Note that the pairs are offset slightly for visibility and that the actual T-D distance is between the 2 points. Significant regression effects across T-D distance and comparisons within eccentricity pairs are indicated with ***: p < 0.001.

Conversely, at *long processing* times T-D distance was only correlated with the sum curvature (R^2^ = 0.0277, F(1,1110) = 31.65, p = 6.06X10^-8^), and not endpoint angle (R^2^ = 0.0005, F(1,880) = 0.432, p = 0.5491). Pairwise comparisons revealed no differences in sum curvature when target and distractor eccentricities varied while the T-D distance was equal (all p > 0.2441), supporting allocentric rather than egocentric effects. There were no effects for saccade endpoint angles when T-D distance was equal across varying target and distractor eccentricities.

## Discussion

In this study, we varied the distance of target and distractor objects relative to a central fixation point, and one another, in a saccadic delayed match to sample task to investigate the reference frame of object-based suppression in the oculomotor system. We found that for short object processing times, while the competition between target and distractor is active^[14]^, both target and distractor are represented in an egocentric reference frame. However, the completion of decision making led to the emergence of distractor suppression, which occurred in an allocentric reference frame.

Saccades executed at short processing times displayed a strong bias for selecting objects closer to fixation (less eccentric). Additionally, the trajectory shifts of these saccades towards the distractor were consistent with weighted vector averaging, as has been previously reported for saccade deviations towards distracting objects^[14–18][21–24][29–30][45]^. Our results for both the saccade trajectories and accuracy suggest that the weight of each object decreases linearly as its eccentricity increases. The strength of this spatial bias alters saccade target selection independent of the identity of the stimuli. Our results lend support to the idea that early saccades are based on bottom-up salience rather than top-down relevance^[53]^ which affects both accuracy and saccade curvature^[54]^. The eccentricity gradient may result from either an inherent attentional bias^[50–52]^, or reflect a visually-based salience difference (i.e. cortical magnification). Both of these possibilities reflect an egocentric reference frame early in the formation of object representations.

In contrast, long processing times prior to saccade execution resulted in deviations away from the distractor indicating that the distractor had been suppressed^[19–20][25–26][29–30][45]^. When the stimuli are processed for longer prior to initiating the saccade, the target is identified and we find that the distractor is suppressed in an allocentric reference frame. The magnitude of the curvature is no longer affected by the eccentricity of the distractor. Instead, the magnitude of the curvature increases as the distance between the target and distractor increases, reflecting a shift from an egocentric to allocentric reference frame. However, there continues to be an effect of *target* eccentricity at long processing times, indicating that an egocentric reference frame is still being utilized. This arises from saccades to targets being programmed in egocentric coordinates^[31–30]^. Object-based suppression in an allocentric reference frame has been found in object-based inhibition of return (IOR)^[34–35]^, and to previous saccade locations^[36–37][55–56]^. The current work extends these findings, identifying resolution of target/distractor competition as the initiator of distractor suppression that occurs in an allocentric frame of reference.

The stimuli used in our study are similar to alphanumeric characters^[40]^, which have previously been suggested to require late-stage ventral stream visual processing in IT cortex^[39–40]^ where objects are represented allocentrically^[41–43]^. The transformation from a gaze-centered reference frame to hand-centered to guide reaching has previously been shown to occur in the posterior parietal cortex (PPC)^[57–58]^, making it the most probable site for task-relevant reference frame transitions(Andersen and Zipser, 1988)^[12][59]^. Therefore, this would suggest that object-based distractor suppression arising in area IT^[60]^ would be carried by projections to PPC^[61]^ subsequently feeding into the oculomotor system (i.e. SC, FEF). As PPC is associated with multiple reference frames^[12][59][62–64]^, it would be able to map allocentric object suppression onto the gaze-centered egocentric reference frame used in saccade planning.

The simultaneous presence of multiple items necessitates discrimination of the target from a distractor. The suppression of the distractor would provide an effective mechanism to improve target selection and motor planning. Our finding that this suppression is allocentric highlights the involvement of object-based distractor inhibition outside of the motor system influencing the saccade planning process. We find that we can measure this allocentric object-based attentional mechanisms in a temporally precise manner manifested through saccade curvatures.

In summary, object-based suppression occurs in an allocentric reference frame affecting target selection and the subsequent motor action. The early representation of potential saccade targets is egocentric and strongly weighted by eccentricity as they compete for selection, with a bias for the object closest to the center of gaze. With longer processing time, the target is identified, and the distractor is suppressed. The target continues to be represented in an egocentric reference frame, while the strength of distractor suppression is dependent on target-distractor distance in an allocentric reference frame. Object-based suppression likely is a general mechanism of visual processing that affects motor planning, as seen here, as well as perceptual decision making, as seen in prior studies.

## Acknowledgements

Funding was provided by NSERC Discovery (RGPIN-2016-05296) and CIHR Operating (102482) grants to MF, and a University of Guelph Undergraduate Summer Research Assistantship to CEO.

## Author contributions

All authors designed the study. CEO collected and analyzed the data. All authors interpreted the results. CEO wrote the draft manuscript. HJ and MF revised the draft. All authors reviewed and approved the final manuscript.

## Data Availability

The data used in this study are being analysed for a second manuscript but can be made available from the authors upon reasonable request. Requests should be directed to the corresponding author.

## Additional Information

The authors declare no competing interests.

## References

1. Ball, K., Smith, D., Ellison, A. & Schenk, T. Both egocentric and allocentric cues support spatial priming in visual search. Neuropsychologia 47, 1585–1591 (2009).

2. Galati, G., Pelle, G., Berthoz, A. & Committeri, G. Multiple reference frames used by the human brain for spatial perception and memory. Experimental brain research 206, 109– 20 (2010).

3. Ruotolo, F. et al. Neural correlates of egocentric and allocentric frames of reference combined with metric and non-metric spatial relations. Neuroscience 409, 235–252 (2019).

4. Burgess, N. Spatial memory: how egocentric and allocentric combine. Trends in Cognitive Sciences 10, 551–557 (2006).

5. Tsotsos, J. K. et al. Modeling visual attention via selective tuning. Artificial Intelligence 78, 507–545 (1995).

6. Desimone, R. & Duncan, J. Neural Mechanisms of Selective Visual Attention. Annu. Rev. Neurosci. 18, 193–222 (1995).

7. Moore, T., Armstrong, K. M. & Fallah, M. Visuomotor Origins of Covert Spatial Attention. Neuron 40, 671–683 (2003).

8. Moore, T. & Fallah, M. Control of eye movements and spatial attention. Proceedings of the National Academy of Sciences 98, 1273–1276 (2001).

9. Moore, T. & Fallah, M. Microstimulation of the Frontal Eye Field and Its Effects on Covert Spatial Attention. Journal of Neurophysiology 91, 152–162 (2004).

10. Balan, P. F. & Ferrera, V. P. Effects of Gaze Shifts on Maintenance of Spatial Memory in Macaque Frontal Eye Field. J. Neurosci. 23, 5446–5454 (2003).

11. Cassanello, C. R. & Ferrera, V. P. Computing vector differences using a gain field-like mechanism in monkey frontal eye field. The Journal of Physiology 582, 647–664 (2007).

12. Andersen, R. A. & Zipser, D. The role of the posterior parietal cortex in coordinate transformations for visual–motor integration. Can. J. Physiol. Pharmacol. 66, 488–501 (1988).

13. Stricanne, B., Andersen, R. A. & Mazzoni, P. Eye-centered, head-centered, and intermediate coding of remembered sound locations in area LIP. Journal of Neurophysiology 76, 2071–2076 (1996).

14. Giuricich, C., Green, R. J., Jordan, H. & Fallah, M. Target–Distractor Competition Modulates Saccade Trajectories in Space and Object Space. eNeuro 10, (2023).

15. Sheliga, B. M., Riggio, L. & Rizzolatti, G. Orienting of attention and eye movements. Exp Brain Res 98, 507–522 (1994).

16. Sheliga, B. M., Riggio, L. & Rizzolatti, G. Spatial attention and eye movements. Exp Brain Res 105, 261–275 (1995).

17. Van der Stigchel, S., Meeter, M. & Theeuwes, J. Eye movement trajectories and what they tell us. Neuroscience & Biobehavioral Reviews 30, 666–679 (2006).

18. Van der Stigchel, S. Recent advances in the study of saccade trajectory deviations. Vision Research 50, 1619–1627 (2010).

19. McSorley, E., Haggard, P. & Walker, R. Time Course of Oculomotor Inhibition Revealed by Saccade Trajectory Modulation. Journal of Neurophysiology 96, 1420–1424 (2006).

20. McSorley, E., Haggard, P. & Walker, R. The spatial and temporal shape of oculomotor inhibition. Vision Research 49, 608–614 (2009).

21. McSorley, E. & McCloy, R. Saccadic eye movements as an index of perceptual decision-making. Exp Brain Res 198, 513–520 (2009).

22. van Zoest, W., Donk, M. & Van der Stigchel, S. Stimulus-salience and the time-course of saccade trajectory deviations. Journal of Vision 12, 16 (2012).

23. Wang, Z. & Theeuwes, J. Distractor Evoked Deviations of Saccade Trajectory Are Modulated by Fixation Activity in the Superior Colliculus: Computational and Behavioral Evidence. PLOS ONE 9, e116382 (2014).

24. Kehoe, D. H., Rahimi, M. & Fallah, M. Perceptual color space representations in the oculomotor system are modulated by surround suppression and biased selection. Frontiers in Systems Neuroscience 12, 1 (2018).

25. Kehoe, D. H. & Fallah, M. Rapid accumulation of inhibition accounts for saccades curved away from distractors. Journal of Neurophysiology 118, 832–844 (2017).

26. Castellotti, S., Szinte, M., Del Viva, M. M. & Montagnini, A. Saccadic trajectories deviate toward or away from optimally informative visual features. iScience 26, 107282 (2023).

27. Robinson, D. A. Eye movements evoked by collicular stimulation in the alert monkey. Vision Research 12, 1795–1808 (1972).

28. Aizawa, H. & Wurtz, R. H. Reversible Inactivation of Monkey Superior Colliculus. I. Curvature of Saccadic Trajectory. Journal of Neurophysiology 79, 2082–2096 (1998).

29. McPeek, R. M., Han, J. H. & Keller, E. L. Competition Between Saccade Goals in the Superior Colliculus Produces Saccade Curvature. Journal of Neurophysiology 89, 2577– 2590 (2003).

30. McPeek, R. M. Incomplete Suppression of Distractor-Related Activity in the Frontal Eye Field Results in Curved Saccades. Journal of Neurophysiology 96, 2699–2711 (2006).

31. Ottes, F. P., Van Gisbergen, J. A. M. & Eggermont, J. J. Visuomotor fields of the superior colliculus: A quantitative model. Vision Research 26, 857–873 (1986).

32. Sommer, M. A. & Wurtz, R. H. Composition and Topographic Organization of Signals Sent From the Frontal Eye Field to the Superior Colliculus. Journal of Neurophysiology 83, 1979–2001 (2000).

33. Ben Hamed, S., Duhamel, J.-R., Bremmer, F. & Graf, W. Representation of the visual field in the lateral intraparietal area of macaque monkeys: a quantitative receptive field analysis. Exp Brain Res 140, 127–144 (2001).

34. Jordan, H. & Tipper, S. P. Object-based inhibition of return in static displays. Psychonomic Bulletin & Review 5, 504–509 (1998).

35. Tipper, S. P., Jordan, H. & Weaver, B. Scene-based and object-centered inhibition of return: Evidence for dual orienting mechanisms. Perception & Psychophysics 61, 50–60 (1999).

36. Sogo, H. & Takeda, Y. Effect of previously fixated locations on saccade trajectory during free visual search. Vision Research 46, 3831–3844 (2006).

37. Pertzov, Y., Zohary, E. & Avidan, G. Rapid Formation of Spatiotopic Representations As Revealed by Inhibition of Return. J. Neurosci. 30, 8882–8887 (2010).

38. Godijn, R., Theeuwes, J. & Godijn, R. The relationship between inhibition of return and saccade trajectory deviations. J Exp Psychol Hum Percept Perform 30, 538–554 (2004).

39. Kehoe, D. H., Lewis, J. & Fallah, M. Oculomotor target selection is mediated by complex objects. Journal of neurophysiology 126, 845–863 (2021).

40. Nobre, A. C., Allison, T. & McCarthy, G. Word recognition in the human inferior temporal lobe. Nature 372, 260–263 (1994).

41. Ito, M., Tamura, H., Fujita, I. & Tanaka, K. Size and position invariance of neuronal responses in monkey inferotemporal cortex. Journal of Neurophysiology 73, 218–226 (1995).

42. Desimone, R. & Gross, C. G. Visual areas in the temporal cortex of the macaque. Brain Research 178, 363–380 (1979).

43. Boussaoud, D., Desimone, R. & Ungerleider, L. G. Visual topography of area TEO in the macaque. Journal of Comparative Neurology 306, 554–575 (1991).

44. Kehoe, D. H., Aybulut, S. & Fallah, M. Higher order, multifeatural object encoding by the oculomotor system. Journal of Neurophysiology 120, 3042–3062 (2018).

45. Kehoe, D. H. & Fallah, M. Oculomotor feature discrimination is cortically mediated. Front Syst Neurosci 17, 1251933 (2023).

46. van Leeuwen, J., Smeets, J. B. J. & Belopolsky, A. V. Forget binning and get SMART: Getting more out of the time-course of response data. Atten Percept Psychophys 81, 2956–2967 (2019).

47. Cnaan, A., Laird, N. M. & Slasor, P. Using the general linear mixed model to analyse unbalanced repeated measures and longitudinal data. Statistics in Medicine 16, 2349– 2380 (1997).

48. Wallace, D. & Green, S. B. Analysis of Repeated Measures Designs with Linear Mixed Models. in Modeling Intraindividual Variability With Repeated Measures Data (Psychology Press, 2001).

49. Krueger, C. & Tian, L. A Comparison of the General Linear Mixed Model and Repeated Measures ANOVA Using a Dataset with Multiple Missing Data Points. Biological Research For Nursing 6, 151–157 (2004).

50. Wolfe, J. M., O’Neill, P. & Bennett, S. C. Why are there eccentricity effects in visual search? Visual and attentional hypotheses. Perception & Psychophysics 60, 140–156 (1998).

51. Van Heusden, E., Olivers, C. N. L. & Donk, M. The effects of eccentricity on attentional capture. Atten Percept Psychophys 86, 422–438 (2024).

52. Yoo, S.-A., Tsotsos, J. K. & Fallah, M. The Attentional Suppressive Surround: Eccentricity, Location-Based and Feature-Based Effects and Interactions. Frontiers in Neuroscience 0, 710 (2018).

53. Fecteau, J. H. & Munoz, D. P. Salience, relevance, and firing: a priority map for target selection. Trends in Cognitive Sciences 10, 382–390 (2006).

54. White, B. J., Theeuwes, J. & Munoz, D. P. Interaction between Visual- and Goal-related Neuronal Signals on the Trajectories of Saccadic Eye Movements. Journal of Cognitive Neuroscience 24, 707–717 (2012).

55. Jonikaitis, D. & Belopolsky, A. V. Target–Distractor Competition in the Oculomotor System Is Spatiotopic. J. Neurosci. 34, 6687–6691 (2014).

56. van Leeuwen, J. & Belopolsky, A. V. Distractor displacements during saccades are reflected in the time-course of saccade curvature. Sci Rep 8, 2469 (2018).

57. Buneo, C. A., Batista, A. P., Jarvis, M. R. & Andersen, R. A. Time-invariant reference frames for parietal reach activity. Exp Brain Res 188, 77–89 (2008).

58. Bremner, L. R. & Andersen, R. A. Temporal Analysis of Reference Frames in Parietal Cortex Area 5d during Reach Planning. Journal of Neuroscience 34, 5273–5284 (2014).

59. Cohen, Y. E. & Andersen, R. A. A common reference frame for movement plans in the posterior parietal cortex. Nat Rev Neurosci 3, 553–562 (2002).

60. Ramezanpour, H. & Fallah, M. The role of temporal cortex in the control of attention. Current Research in Neurobiology 3, 100038 (2022).

61. Choi, S. H., Jeong, G., Kim, Y. B. & Cho, Z. H. Proposal for human visual pathway in the extrastriate cortex by fiber tracking method using diffusion-weighted MRI. NeuroImage 220, 117145 (2020).

62. Zaehle, T. et al. The neural basis of the egocentric and allocentric spatial frame of reference. Brain Research 1137, 92–103 (2007).

63. Chen, Z. Object-based attention: A tutorial review. Atten Percept Psychophys 74, 784– 802 (2012).

64. Derbie, A. Y. et al. Common and distinct neural trends of allocentric and egocentric spatial coding: An ALE meta-analysis. European Journal of Neuroscience 53, 3672–3687 (2021).

